# VPHYL: A Web Application for Converting Emu and EPI2ME wf-16s Nanopore 16S rRNA Abundance Tables into phyloseq and BIOM Objects

**DOI:** 10.64898/2026.11.18.767361

**Authors:** Venkata Sai Reddy Ramireddy, Sarvesh Kumar Srivastava

**Affiliations:** Centre for Biomedical Engineering, Indian Institute of Technology Delhi, New Delhi, India; Department of Biomedical Engineering, All India Institute of Medical Sciences, New Delhi, India

## Abstract

Taxonomic abundance data derived from 16S rRNA gene sequencing data are fundamental in microbiome research. To perform downstream statistical analysis and visualization, raw abundance data must typically be converted into standardized data structures, such as phyloseq S4 objects or BIOM files. This conversion requires three input data files: an Operational Taxonomic Unit (OTU) table (count of specific taxa across samples), a Taxonomy table (hierarchical lineage from Kingdom to Species), and a Sample Data table (experimental metadata). For Oxford Nanopore (ONT) 16S data, the two commonly used classifiers, Emu and the EPI2ME wf-16s workflow (Minimap2/Kraken2), generate output files with different formats and column schemas. We found no web-based tool that accepts these classifier outputs directly. Experienced users can reshape them with a script. Researchers who do not program cannot, and for them, this step remains a hurdle. We introduce VPHYL(Versatile PHYLogenetic Standardizer), an R Shiny web application that accepts abundance outputs directly from Emu and EPI2ME wf-16s, splits the lineage into standard ranks, attaches sample metadata, and generates phyloseq objects or BIOM files. VPHYL integrates multi-file aggregation and produces automated HTML reports comprising read statistics, alpha diversity metrics, and taxonomic abundance visualizations. VPHYL output was identical to that of an independently written R script for all nine supported classifier-database combinations. We validated the framework by processing raw abundance datasets processed using the EPI2ME Labs 16S Workflow from a published respiratory microbiome study comprising both porcine and human microbiota composition samples. The published tables were processed by VPHYL without modification, and the resulting alpha diversity indices and taxonomic profiles were consistent with the reported findings; small differences reflect read filtering applied in the original study.

The application is publicly available at https://venkat-247.shinyapps.io/VPHYL/.

## 1. Introduction

Amplicon sequencing has become a fundamental approach for profiling microbial communities across diverse environments to characterize community composition, diversity, and ecological dynamics. Amplicon sequencing of the 16S rRNA gene is used to study microbial communities across environments, from human health [1] to oceans [2]. Short-read sequencing platforms like Illumina have dominated the sequencing of the 16S subunit of the ribosomal RNA gene because of its highly per-base accuracy, but short-read methods are restricted by read-length constraints (∼300 bp), limiting sequencing to isolated hypervariable regions (V3-V4), restricting the taxonomic resolution to genus level.[3]In this context, long-read 16S rRNA sequencing via Oxford Nanopore Technology (ONT) has become a preferred approach in microbiome research due to its affordability and ability to sequence all nine hypervariable regions(V1-V9) to achieve species-level resolution. [4]However, the limitation of the ONT is that it has a high rate of sequencing per-base errors. This makes it difficult to distinguish between closely related species that share high sequence similarity. [5], [6].PacBio HiFi (CCS) sequencing also provides full-length 16S reads with higher per-base accuracy than ONT[7]. VPHYL addresses ONT outputs only

The high error profile of Nanopore amplicon data has led to the rise of multiple classifiers, which are Centrifuge, Nanoclust and MetaMaps[8], [9], [10]. While numerous tools exist, two pipelines are frequently used for the ONT 16S ecosystem due to accessibility and accuracy: the EPI2ME 16S workflow and Emu (see Supplementary Information **Figure 1** for an overview of the end-to-end analytical workflow). EPI2ME is the official, vendor-supported cloud and local workflow for Nanopore users, relying on Kraken2 (rapid, exact k-mer matching) and Minimap2 (error-tolerant seed-chain-align strategies). Emu was developed specifically for error- prone Nanopore 16S amplicon data. Emu utilizes an Expectation-Maximization (EM) algorithm to iteratively evaluate the mapping density of all reads in a sample, probabilistically resolving ambiguous assignments to deliver highly accurate species-level quantification that minimizes the false-positive rates seen in k-mer approaches [11], [12]. The accuracy of classifiers depends on the design and curation standards of the reference databases. Major reference databases, including the Ribosomal Database Project (RDP) is based on a Naïve Bayesian framework for assigning taxonomy to partial reads [13]. The SILVA 138.1 database is widely used for diversity studies due to its manually curated subunit alignments [14], [15]. The NCBI targeted loci (16S/18S/28S/ITS) offer a comprehensive nucleotide database [16]. Specialized indices like the Emu Default Database integrate high-quality RefSeq data with the Ribosomal RNA Database (rrnDB) metrics to correct for 16S gene copy-number bias [11], [17]. Because no single database provides a perfect balance of taxonomic resolution, phylogenetic accuracy, and nomenclature standards, the choice of the database depends on the experimental objectives.

**Figure 1:**
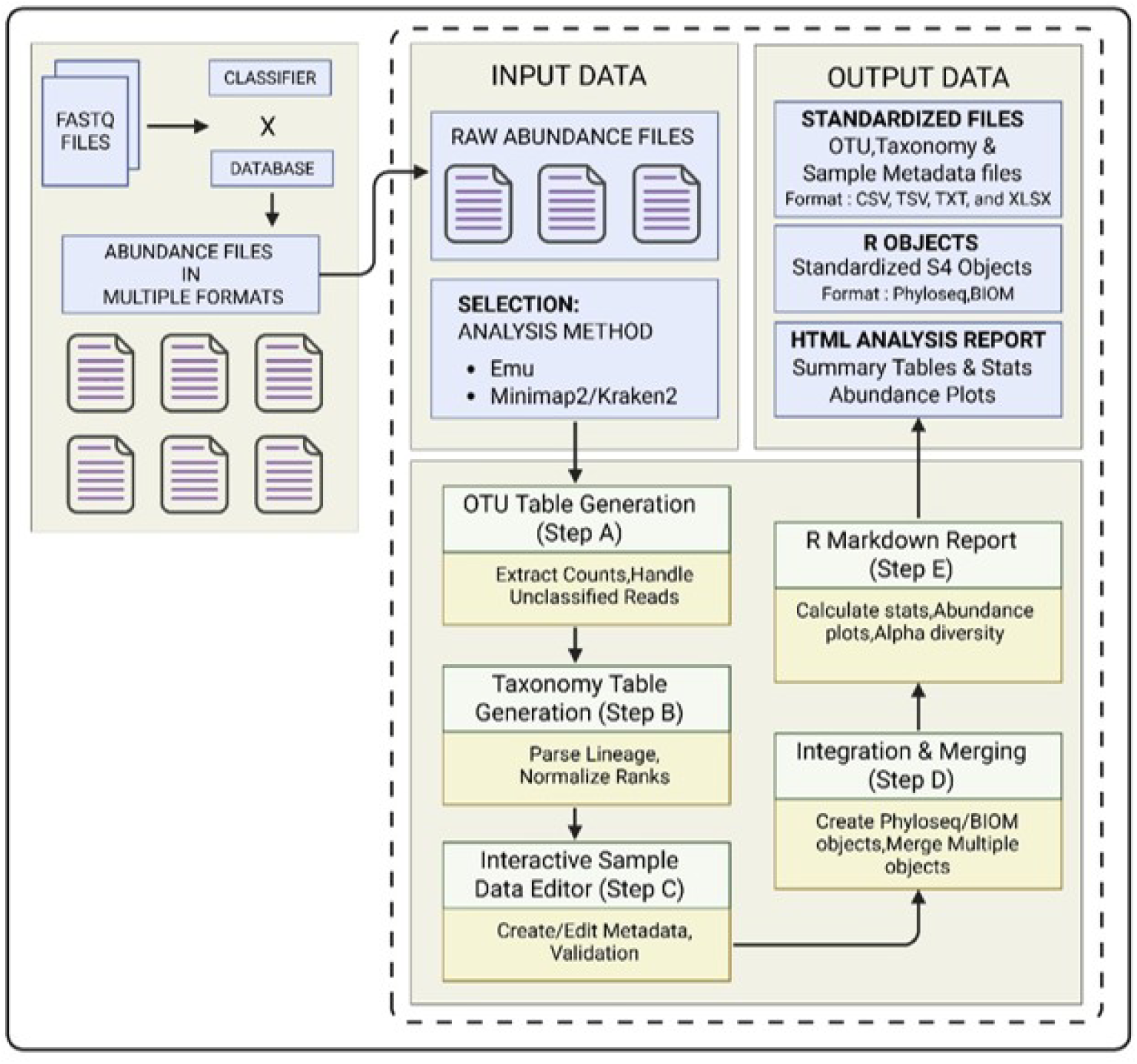
VPHYL: Graphical summary of graphical interface and framework for an automated method for standardized multi-format output from Emu, EPI2ME 16S Workflow (Minimap2, and Kraken2) pipelines into downstream-compatible R objects.

This use of classifier-database combinations, such as the EPI2ME workflow (utilizing Kraken2/Minimap2 against NCBI and SILVA) and the Emu pipeline (with Emu- default, RDP, or SILVA), introduces data heterogeneity. Each pairing generates an abundance table with its own column layout, headers and taxonomic naming. For example, a SILVA-based Emu run gives a single semicolon-delimited lineage column, while an NCBI-based Kraken2 run generates separate rank-specific headers with different taxonomic terminology (e.g., ‘superkingdom’ vs. ‘Kingdom’). Converting these tables into the count matrix, taxonomy matrix and sample metadata that phyloseq[18] or a BIOM table requires is a short scripting task for users fluent in R or Python. Emu’s combine-outputs subcommand already merges per-sample outputs into an abundance table and, optionally, a taxonomy table, and a delimited lineage can be split in a few lines of code. Researchers who do not script, however, still need such a file before they can use no-code platforms that accept BIOM input, such as MicrobiomeAnalyst [19], or pass their data to an analyst (Supplementary Information Figure 1). Several tools address parts of this problem, mostly as command-line programs or R packages. The epi2me2r R package imports WIMP and antimicrobial- resistance CSV files from the EPI2ME platform into phyloseq or metagenomeSeq objects. Ecosystem-specific scripts such as kraken-biom and the KrakenTools suite successfully convert Kraken2-generated reports into downstream-ready formats, but they remain strictly constrained to the Kraken architecture and cannot parse Emu or EPI2ME outputs [20], [21]. Recently, TAXPASTA (TAXonomic Profile Aggregation and STAndardisation) has been developed as a Python library that aggregates outputs from metagenomic profilers like Bracken, Centrifuge, DIAMOND, ganon, Kaiju, Kraken2, KrakenUniq, MALT/MEGAN6, MetaPhlAn, and mOTUs into standardized tabular formats [22]. Emu is not among the profilers TAXPASTA supports. A recent attempt to solve the interoperability of microbiome data is the Core Analysis Modular Pipeline (CAMP), a modular, Snakemake-based metagenomics analysis system that accepts the multi-format output generated by different classifiers like MetaPhlan4, Kraken2/Bracken, and/or XTree. CAMP attempts to solve the problem by running MetaPhlAn4, Kraken2/Bracken, and XTree internally on the same raw reads and then normalizing their divergent output formats into a common downstream-ready table within the same pipeline execution. CAMP begins from raw sequencing reads and runs on the command line, so it serves a different stage of the workflow than a precomputed Emu or wf-16s table[23]. QIIME 2 can import externally generated tables in BIOM format, but this requires the command line and a pre-formatted BIOM file [24], [25]. MicrobiomeAnalyst provides end-to-end processing from raw reads and also accepts count, taxonomy and metadata tables or a BIOM file. At the same time, Pavian visualises Kraken-style reports, and neither reads Emu nor wf-16s tables directly [19], [26].

To address these limitations, we developed VPHYL(Versatile PHYLogenetic Standardizer), an interactive R Shiny-based web platform for Nanopore 16S abundance tables. VPHYL accepts already-classified Emu or EPI2ME abundance tables directly, without requiring command-line use or re-classification from raw reads and converts them deterministically into standard phyloseq or BIOM objects. VPHYL makes three contributions. First, it checks the headers of an uploaded table against the method selected by the user and applies the corresponding parsing rule. Second, it standardizes heterogeneous lineage strings and rank-specific headers from Emu and EPI2ME output into a single seven-level taxonomy using a fixed heuristic rule set, so the same input always produces the same output. Third, it exposes this conversion through a browser-based interface, so a user without R or command-line experience can move from a raw classifier output to a phyloseq or BIOM object ready for downstream analysis. The current implementation is scoped to Emu and EPI2ME wf-16s outputs from ONT 16S data. VPHYL makes this step reproducible for users who do not script.

## 2. Methods

### 2.1. Implementation Environment: R Shiny Framework

The VPHYL application is developed using the Shiny framework. [27] within the R statistical environment [28]. Shiny was selected as the development platform to bridge the gap between complex command-line bioinformatics pipelines and end- users lacking extensive programming experience. By using Shiny’s reactive programming model, VPHYL used the data structures of the phyloseq package [18] behind a graphical user interface (GUI) (**Figure 1**). This approach allows for the real- time manipulation of taxonomic matrices and sample metadata without requiring the user to manually execute custom scripts. Supplementary Information **Table 1** provides a detailed comparison of VPHYL with other tools in the space of microbiome analysis.

**Table 1:** Detailed breakdown of Alpha diversity indices (Shannon and Observed OTUs) comparing the re-analysis performed by the VPHYL platform against the published Nanopore sequencing results from Macip et al.. The comparison spans both the Human and porcine respiratory microbiome datasets to validate the platform.

| **Dataset** | **Index** | **VPHYL Platform** | **Macip et al. 2025** |
|---|---|---|---|
| Porcine | Shannon Index Range | 0.88 - 2.33 | 0.80 - 2.29 |
|  | Observed OTUs Range | 16 - 89 | 22 - 89 |
|  | Mean Shannon | 1.45 | 1.474 |
|  | Mean Observed OTUs | 63.4 | 64.29 |
| Human | Shannon Index Range | 0.104 - 2.749 | 0.10 - 2.75 |
|  | Observed OTUs Range | 4-56 | 4-51 |
|  | Mean Shannon | 1.56 | 1.504 |
|  | Mean Observed OTUs | 23.1 | 20.75 |

### 2.2. Application Description

VPHYL handles the step after taxonomic classification of Nanopore 16S data. Its primary function is the automated parsing, normalization, and integration of heterogeneous taxonomic abundance data generated from the classifier-database combinations of EPI2ME inbuilt 16S workflow pipelines (Kraken2 and Minimap2 mapped against NCBI_16s_18s, NCBI_16s_18s_28s_ITS, and SILVA_138_1 databases) and the Emu pipeline (utilizing Emu_Default, RDP_v11.5, and SILVA_v138.1 databases) into S4 class standardized R objects. The tool automates the decomposition of raw abundance files into Operational Taxonomic Unit (OTU) tables, taxonomy matrices, and sample metadata, subsequently merging these components into a unified object suitable for downstream and microbial analysis (**Figure 1**)

To ensure the validity of the file uploaded and the choice of method selected, VPHYL checks the column headers upon file upload. The system scans the column headers of the input file to confirm that they match the selected method. For the EMU method, the algorithm validates the presence of a *tax_id* identifier and associated taxonomic lineage columns (e.g., *tax_id,* genus, species). Similarly, for EPI2ME workflows, the system scans for the concatenation of taxonomic ranks within a single *tax* column. This automated detection prevents the format mismatches that cause compilation errors in manual workflows. The interface guides the user through four steps (A-D); each step unlocks only after the previous one is validated. The UI also features real-time feedback by providing the user status update process (Supplementary Information **Figures 2** and 3).

**Figure 2:**
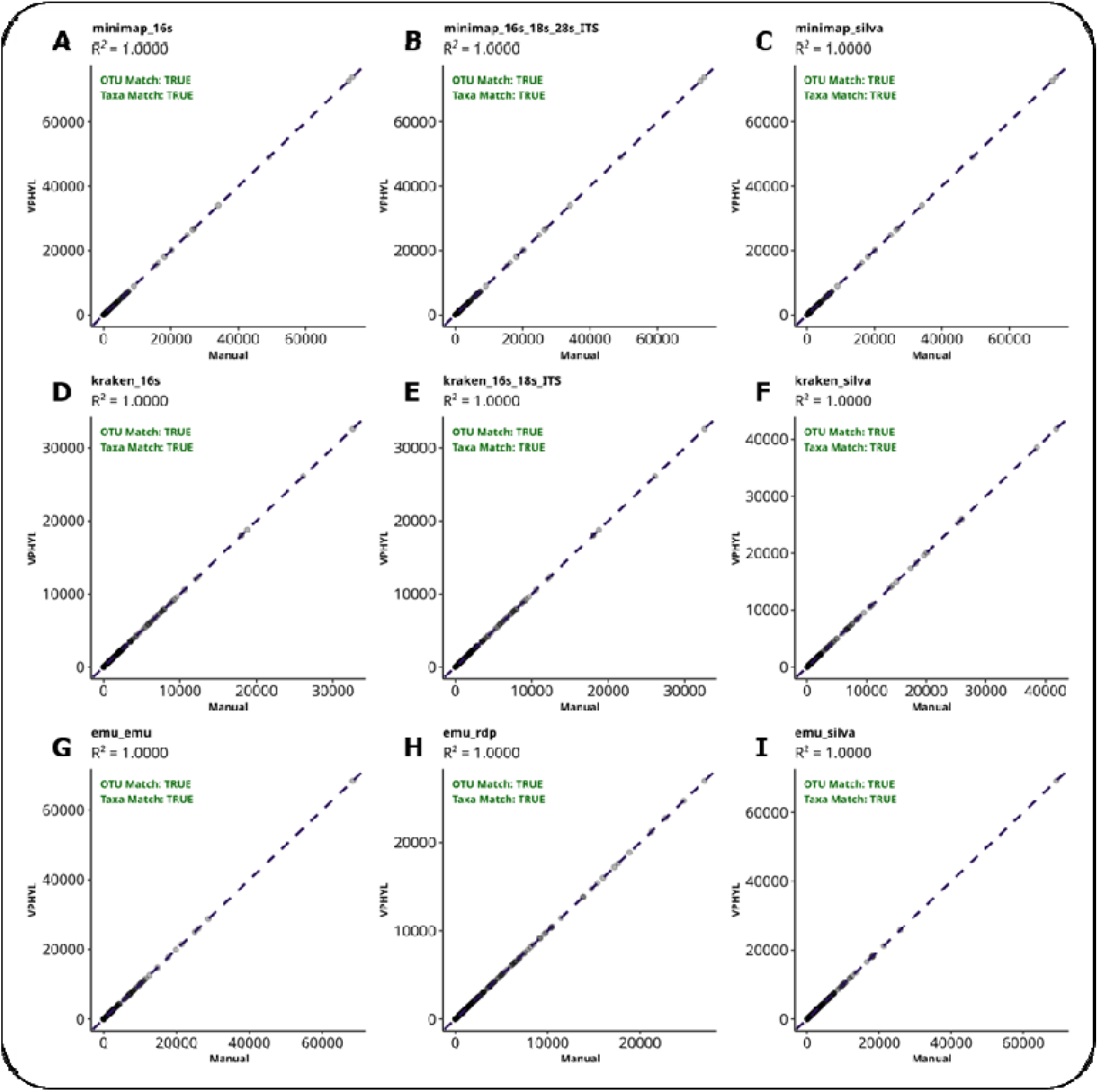
Validation of Taxonomic Abundance Preservation Between Manual Parsing and VPHYL Processing. Scatter plots comparing taxon abundance counts derived from manually parsed baseline datasets (x-axis) and VPHYL-processed datasets (y-axis) to verify that automated data ingestion preserves taxonomic abundance information. All workflows showed perfect agreement (R² = 1.0000), with complete OTU and taxa matching between manually processed and VPHYL- generated datasets. Panels A-C are outputs from the EPI2ME 16S workflow utilizing the Minimap2 classifier against the (A) NCBI 16 S, (B) NCBI 16S/18S/28S/ITS, and (C) SILVA v138.1 databases. Panels D-F are outputs from the EPI2ME 16S workflow utilizing the Kraken2 classifier against the (D) NCBI 16S, (E) NCBI 16S/18S/ITS, and (F) SILVA v138.1 databases. Panels G-I are outputs from the Emu classifier paired with the (G) Emu default, (H) RDP, and (I) SILVA databases.

**Figure 3:**
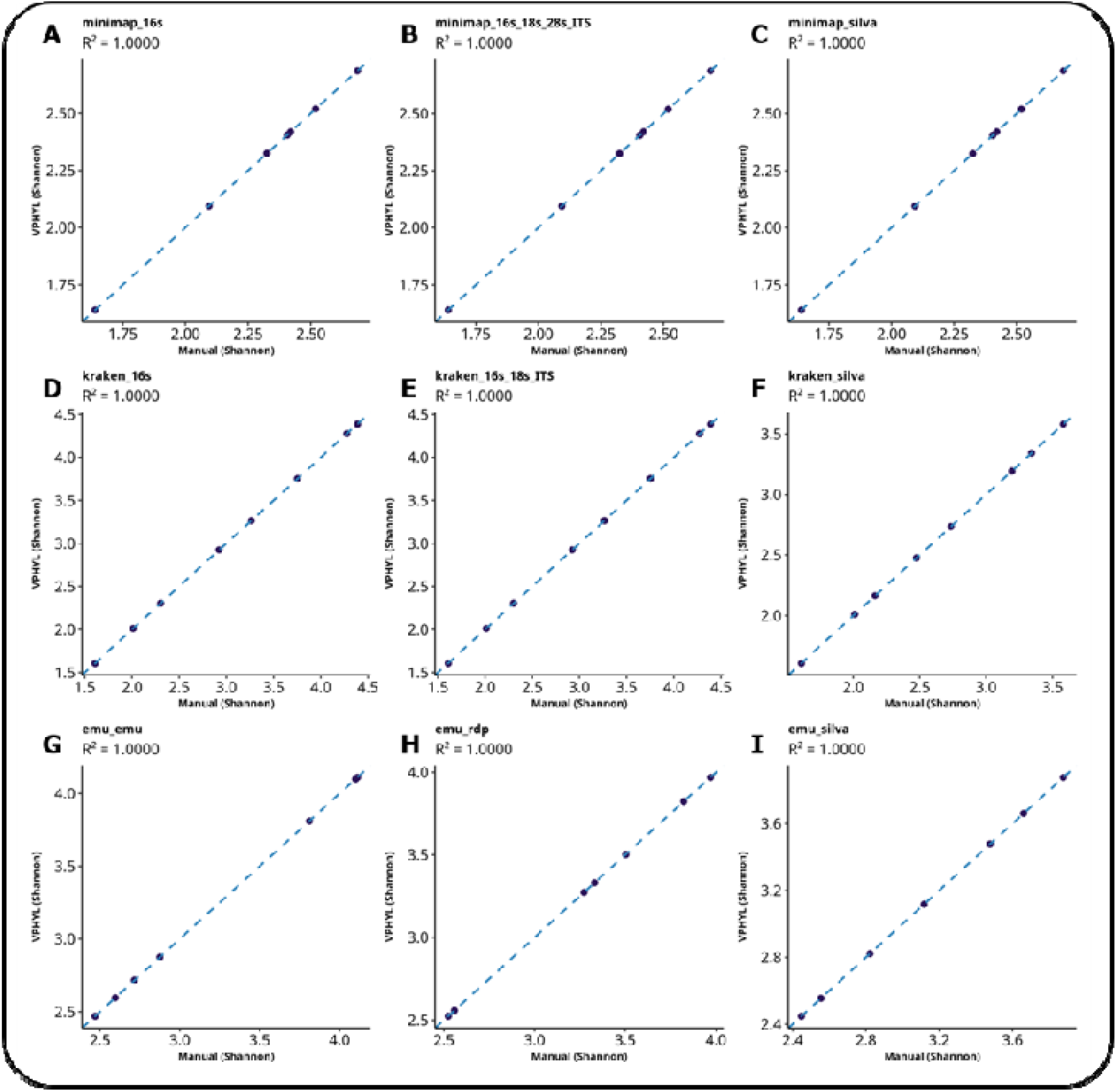
Alpha Diversity Assessed via Within-Sample Diversity Metrics. Scatter plots illustrating the correlation of calculated alpha diversity (Shannon) metric derived from manually parsed baseline datasets (x-axis) and VPHYL-processed datasets (y- axis) to verify that automated data ingestion does not alter alpha diversity. Panels A- C are outputs from the EPI2ME 16S workflow utilizing the Minimap2 classifier against the (A) NCBI 16S, (B) NCBI 16S/18S/28S/ITS, and (C) SILVA v138.1 databases. Panels D-F are outputs from the EPI2ME 16S workflow utilizing the Kraken2 classifier against the (D) NCBI 16S, (E) NCBI 16S/18S/ITS, and (F) SILVA v138.1 databases. Panels G-I are outputs from the Emu classifier paired with the (G) Emu default, (H) RDP, and (I) SILVA databases.

### 2.3. Processing Algorithm and Framework

The core framework of VPHYL is a customised parsing algorithm for generating the OTU and Taxonomy matrices. For Emu Outputs: The algorithm implements a pre- processing method to handle unclassified reads, which are often discarded in standard conversions. The system identifies rows flagged as unmapped or mapped_unclassified in the *tax_id* column and propagates these labels across all taxonomic rank columns. This ensures that unclassified reads are retained in the final abundance calculations and OTU Matrix generation. For taxonomy generation, the system supports both separate-column formats (NCBI/RDP style) and semi- colon-delimited lineage strings (SILVA style), automatically splitting the latter into standard ranks (Kingdom to Species). For EPI2ME Outputs: The parsing logic executes a string-split operation on the *tax* column, utilizing the semi-colon delimiter to populate a temporary taxonomic dataframe. Simultaneously, the abundance data is isolated to form the OTU matrix. To maintain matrix synchronization, row names are forced to be unique identifiers, resolving potential duplicates that occur when multiple OTUs map to the same taxonomic annotation.

A frequent bottleneck in constructing phyloseq objects is the mismatch between sample IDs in the OTU table and the metadata file. VPHYL pre-fills the first column of the metadata table with the sample IDs taken from the abundance matrix, so the two always match. The interactive grid allows the users to manually annotate the samples columns and rows directly within the browser or the user can upload the metadata file. Once the component matrices (OTU, Taxa, and Metadata) are finalized, the system invokes the *phyloseq()* constructor to instantiate the S4 object. Users can upload multiple abundance files sequentially; the system processes them individually and then utilizes the *merge_phyloseq()* function to combine selected objects (Supplementary Information **Figure 4**). This allows samples classified in separate runs to be combined. To provide immediate analytical insight, VPHYL integrates an R Markdown templating engine. Upon request, the system passes the final phyloseq object to a parameterized. Rmd template, which is rendered into a self-contained HTML document. This report utilises ggplot2 [29] and DT [30] To visualize, **i**) Read Statistics: A tabulation of classified versus unclassified reads per sample, **ii**) Abundance Stats: Aggregated counts at Phylum and Species taxonomic level, **iii**) Alpha Diversity: Table containing Observed, Chao1, Shannon and Simpson, **iv**) Compositional visualization: Stacked bar charts displaying the relative abundance of the top 20 taxa at Species and Phylum level.

**Figure 4:**
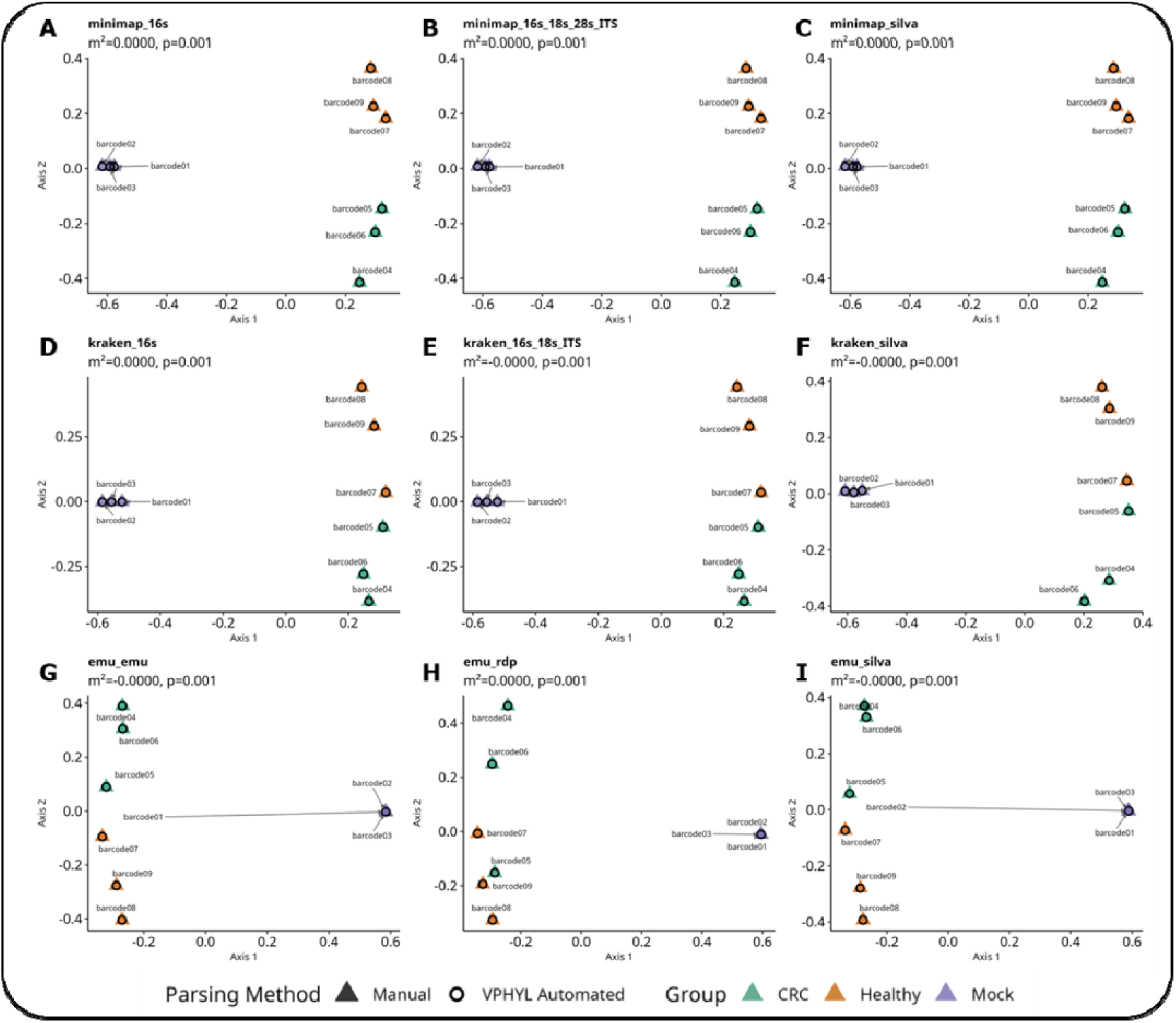
Beta Diversity Assessed via Procrustes Analysis. Procrustes analysis applied to Principal Coordinate Analysis (PCoA) ordinations to measure the similarity, verifying that the data objects automatically generated by VPHYL are structurally equivalent to those derived from manual baseline parsing. PCoA was computed using Bray-Curtis dissimilarity on relative taxon abundance profiles. Panels A-C are outputs from the EPI2ME 16S workflow utilizing the Minimap2 classifier against the **A)** NCBI 16S, **B)** NCBI 16S/18S/28S/ITS, and **C)** SILVA v138.1 databases. Panels D-F are outputs from the EPI2ME 16S workflow utilizing the Kraken2 classifier against the **D)** NCBI 16S, **E)** NCBI 16S/18S/ITS, and **F)** SILVA v138.1 databases. Panels G-I are outputs from the Emu classifier paired with the **G)** Emu default, **H)** RDP, and **I)** SILVA database.

### 2.4. Input Data

VPHYL accepts raw abundance flat files generated directly by the supported classifier-database combinations. Supported formats include TSV, CSV, TXT, and XLSX. The application currently supports Emu outputs processed against the Emu_Default, RDP_v11.5, and SILVA_v138.1 databases, as well as EPI2ME 16S Workflow (Kraken2 or Minimap2) outputs processed against the NCBI_16s_18s, NCBI_16s_18s_28s_ITS, and SILVA_138_1 databases. Users can upload single or multiple abundance files sequentially for their selected analysis method. To assist users in formatting their data and to demonstrate the tool’s capabilities, VPHYL includes a “Load Example Data” feature that populates the application with correctly formatted outputs.

### 2.5. Load Example Data

This test dataset was compiled using a combination of synthetic positive-control samples and real-world clinical sequencing data containing barcode 01-09. The synthetic component has three barcodes (barcodes 01-03) constructed with 17 taxa at defined relative proportions sourced from the ZymoBIOMICS® Gut Microbiome Standard (D6331, Zymo Research). The clinical component contains full-length 16S rRNA gene amplicon sequencing data (V1–V9 regions, sequenced via Oxford Nanopore MinION) sourced from a published colorectal cancer (CRC) study (NCBI BioProject Accession: PRJNA911189). The clinical subset includes fecal samples from both diagnosed CRC patients (barcode 04:CCR-102-1-C, barcode 05:CCR- 103-1-C, barcode 06:CCR-109-1-C) and healthy non-CRC controls (barcode 07:SANO-104-1-C, barcode 08:SANO-105-1-C, barcode 09:SANO-103-1-C).

### 2.6. Web Interface Implementation

The VPHYL user interface is built using Bootstrap 4 themes via the bslib package. The interface is characterised by its dynamic UI elements using ShinyR. Key features of the application and user interface include: **i**) panels for processing (Step A-D) remain hidden until a valid file is loaded, reducing visual clutter. **ii**) Error handling is managed via R’s tryCatch mechanism, which intercepts parsing failures and displays user-friendly notifications. iii) Generate and download OTU, Taxa and sample metadata files in multiple formats. **iv**) Generate and download R-compatible objects for microbiome analysis and sample HTML analysis report.

### 2.7. Output Data

Three types of outputs are generated by VPHYL: i) The processed OTU table, Taxonomy table, and Sample Metadata are downloadable as .txt,.csv, .tsv, or .xlsx files. ii) The final microbiome data structure is exportable as an.RData file (for direct loading into R environments) or a .biom file. iii) The basic summary file is exported as a standalone HTML file, which can be viewed in any web browser without dependency on R or the Shiny server (**Figure 1**).

### 2.8. Performance

VPHYL web application has been tested on standard personal computer (Windows 10: AMD Ryzen 5 4600H with Radeon Graphics (3.00 GHz), 8GB RAM) and desktop (Ubuntu 24.04.2 LTS: Intel® Core™ i9-14900K x32,128GB RAM) in the latest versions of Chrome, Edge, Firefox and Brave browsers. For smaller input files (6-12 KB, approx. 90 rows) generated by Emu, EPI2ME 16S workflow (Minimap2, or Kraken2), the web-hosted application (shinyapps.io) required 5-10 seconds. For a larger dataset (478 KB, 2,029 rows), the hosted application completed processing in 20-25 seconds. When processing the same file locally with RStudio (v4.3.3), the larger 478 KB file was processed in approximately 5 seconds on a high-performance workstation, compared to 40-45 seconds on a standard personal computer.

### 2.9. Limitations

Currently, VPHYL parsing logic relies on specific column headers (e.g., *tax_id*, *tax*) and strictly supports abundance files generated using the Emu tool or the EPI2ME 16S Workflow. The application is hosted on Shinyapps and made accessible through the web, with a maximum file upload limit of 100 MB and a default memory allocation of 1 GB. Analysis of files larger than 10 MB may require increased processing time, leading to longer wait times for the generation of the final HTML report. It has been tested only on ONT 16S data, and taxonomic classification must be completed before using VPHYL. VPHYL converts tables as provided and applies no read or abundance filtering, so values may differ from studies that filtered their data.

## 3. Applications

VPHYL could help the researchers reduce the combinatorial load of parsing and forming heterogeneous raw abundance files while trying to decide the optimal pair of Taxonomic Classifier and Databases. It provides users and researchers a streamlined, reproducible framework to standardize heterogeneous taxonomic abundance data from Nanopore classification pipelines such as Emu and EPI2ME 16S Workflow (Minimap2/Kraken2) that can be automatically processed, standardized into a single unified phyloseq S4 object or BIOM file and downstream- compatible formats to bridge the gap between upstream taxonomic classification and downstream analysis.

## 4. Results

A critical requirement for any automated data-ingestion tool is that it introduces no computational artifacts relative to established or manual workflows, as differences in data processing pipelines have been shown to alter downstream biological conclusions, including the number and identity of significant taxa, alpha diversity estimates, and community structure [31], [32]To evaluate this for VPHYL, we used a multi-level validation that tests equivalence at three analytical scales: (i) element- wise matrix fidelity that verifies individual taxon abundance count and taxonomic annotation is preserved exactly (ii) within-sample summary metrics (alpha diversity) and (iii) between-sample community structure (beta diversity)that establishes the multivariate spatial relationships among samples are identical between workflows. The audit baseline against which VPHYL was compared consisted of a standalone R script that independently reimplements the VPHYL parsing specification including unclassified-read propagation, semicolon-delimited lineage splitting, and rank- nomenclature standardization, but was written and executed independently of the application source code. Both implementations processed a common test dataset comprising nine barcodes: Load Example Data, classified through all nine classifier- database combinations supported by VPHYL.

The most important test of parsing correctness is whether every individual taxon abundance value and its associated taxonomic annotation are preserved during automated ingestion. Errors at this level, such as incorrect rank assignments during lineage splitting can propagate through all downstream analyses. To detect such errors, we performed element-wise comparison of the OTU abundance matrices and taxonomy annotation tables generated by VPHYL against those produced by the independent audit script. Across all nine classifier-database combinations, OTU and taxonomy matrices produced by VPHYL were identical to the audit reference. The taxonomic abundance matrices and taxonomic annotations produced via VPHYL were identical to those obtained via manual parsing for all classifier-database combinations and matched with an R² of 1 (**Figure 2**), showing that VPHYL’s schema-detection logic and lineage-splitting algorithms preserve every individual data element without introducing artifacts.

While matrix identity guarantees that the raw inputs to ecological calculations are correct, it is essential to verify that the derived diversity metrics which are the primary statistical endpoints in most microbiome studies are also preserved. Alpha diversity indices are particularly sensitive to data-handling decisions such as the treatment of unclassified reads, rounding of fractional abundances, and the inclusion or exclusion of low-abundance taxa. [33]Even numerically minor parsing errors can alter the resulting diversity estimates. We therefore computed the Shannon diversity index for all nine samples within each of the nine classifier-database datasets, using both VPHYL-generated and audit-derived phyloseq objects. The Shannon index yielded R² of 1 between VPHYL and audit outputs across every dataset tested (**Figure 3**). This confirms that VPHYL’s handling of unclassified reads and abundance-count rounding preserves the ecological signal at the within-sample level.

The most stringent test of data-processing equivalence is whether the multivariate spatial relationships among all samples are preserved, and whether the relative positions of samples in ordination space remain identical between the two workflows. Beta diversity analyses are affected by systematic biases like subtle changes in the relative proportions of rare community members that may not be visible at the single- metric level. [34]. To assess this we measured the Bray-Curtis dissimilarity matrices from relative taxon abundance profiles for both VPHYL-generated and audit-derived phyloseq objects, projected them into Principal Coordinate Analysis (PCoA) ordination space, and applied Procrustes rotation analysis (vegan::protest(); 999 permutations)[35] to quantify the structural similarity between the two ordination configurations. The Procrustes m² statistic measures the sum of squared residuals after optimal rotation, translation, and scaling of one ordination onto the other(m² = 0 indicates geometric identity between the two configurations)[36] Across all nine classifier-database combinations, the Procrustes analysis returned m² = 0 with a p- value (p < 0.001) in every case (**Figure 4**). This result shows that VPHYL’s automated parsing preserves individual data elements and the full multivariate community structure at the ordination level.

The results from the three validation levels show that VPHYL functions as a deterministic conversion of raw classifier output into structured phyloseq objects. The identical R² = 1.0000 at the matrix level (**Figure 2**), the perfect agreement of the Shannon alpha diversity index (**Figure 3**), and the Procrustes residuals to m² = 0 at the ordination level (**Figure 4**) collectively confirm that no information is lost, altered, or introduced during VPHYL’s automated schema detection, lineage parsing, and matrix construction across all nine supported classifier-database configurations.

## 5. Validation study

To show the effective functionality of the VPHYL platform, we present a comparison of the alpha indices and abundance plots. This comparison is consistent with the findings reported in the study by Macip, Guillem, et al.[3] with the values obtained by processing the raw abundance files included in the supplementary(Supplementary Information **Table 2**,Supplementary Information **Table 3**), utilizing the VPHYL platform hosted on Shiny Apps. The raw abundance files of the above study, from porcine and human samples originally processed with the EPI2ME wf-16s workflow, were uploaded to VPHYL as provided (**Table 1**). The dataset included multiple sequencing time points (12, 18, 24, and 72 hours). The pipeline computed alpha diversity metrics, specifically the Shannon, Simpson, Chao1, and Observed OTU indices, which were cross-referenced with the ranges reported in the original study. Additionally, taxonomic abundance profiles were generated to validate patterns in community composition. In the porcine dataset, Shannon indices ranged from 0.9 to 2.29 across all 10 barcodes, while the observed OTUs varied from 22 to 89 (Supplementary Information **Table T2**). Similarly, in the human dataset, the Shannon indices ranged from 0.104 to 2.749, with observed OTU counts ranging from 4 to 56 (Supplementary Information **Table T3**), aligning with those reported in the paper. The VPHYL-derived values were consistent with the published results; small differences were observed in the “Observed OTUs” metric, particularly in the human dataset (Mean: 23.1 vs. 20.75). These slight discrepancies are attributable to differences in bioinformatic filtering strategies (**Table 1**).

The porcine study dataset (**Figure 5A**) was heavily dominated by a few specific genera. Pseudomonas, Escherichia-Shigella, and Veillonella were the main taxa across the samples, but their relative abundances varied between the different barcodes, and a large portion of the total read counts fell into the “Unclassified” category, suggesting that there’s still a significant part of the porcine respiratory microbiome that is not mapped out. On the other hand, the Human microbiota study dataset (**Figure 5B**) showed a diverse set of taxa, as compared with the porcine dataset. In this case, Pseudomonas and Stenotrophomonas were the most common genera, especially in the higher-yield barcodes, while Escherichia-Shigella, Haemophilus, and Staphylococcus were also dominant taxa in other barcodes. Unlike the porcine data, which showcased consistent dominant genera, the human samples exhibited much more variation among the barcodes, matching the patterns reported in the paper. The main purpose of these validation results is to provide a case study to demonstrate the capability of VPHYL to generate the RData and BIOM files and perform basic microbial visualization and analysis. The variation in the alpha diversity metrics reported in **Table 1** between the reported values and the VPHYL-generated values is because of the filters applied in the paper, as those exact filters were absent in the VPHYL platform. This case study shows that published wf-16s abundance tables can be processed by VPHYL as provided, without programming.

**Figure 5:**
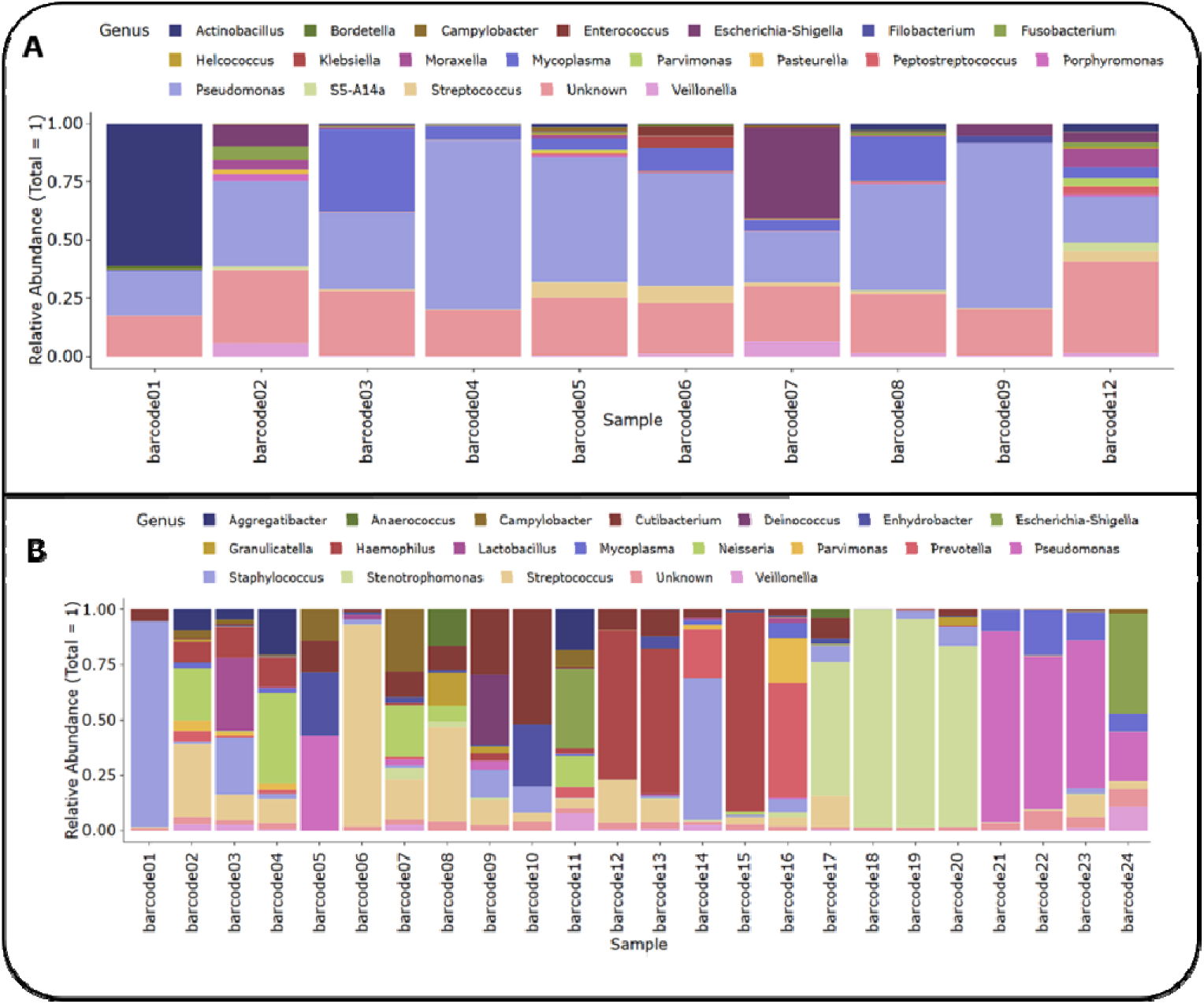
The top 20 abundance plots at the genus level for **A)** Porcine and **B)** Human respiratory microbiome data at 72-hour sequencing duration were created by processing the raw abundance tables from the study by Macip, Guillem, et al [3]. using VPHYL.

## 6. Summary

In summary, Nanopore amplicon sequencing is increasingly adopted for high- resolution microbiome profiling. Yet, the output formats of Emu and wf-16s differ and must be converted before downstream tools can read them. Previous efforts in the field have addressed several dimensions of this challenge: the BIOM format standardized file exchange within the QIIME-era ecosystem to facilitate interoperability across tools and meta-analyses, phyloseq and MicrobiomeAnalyst accept structured input tables, and MicrobiomeAnalyst also offers end-to-end processing from raw reads, and CAMP, TAXPASTA made meaningful progress by accepting multi-format outputs from different classifiers and normalizing them into a common downstream-ready table within a single pipeline execution. We found no tool that accepts the native outputs of both Emu and wf-16s and returns phyloseq and BIOM objects without programming. Researchers fluent in R can script the same conversion, for example with Emu’s combine-outputs subcommand and a few lines of code.VPHYL’s contribution is practical, offering a fixed, documented conversion that needs no programming Future development of VPHYL will prioritise expanding parsing compatibility to additional Nanopore classifiers as they emerge, including tools that integrate custom reference databases or produce non-standard taxonomic rank schemas. Improvements to server-side resource allocation will address the current RAM- dependent processing ceiling for large files, allowing larger files to be processed. Furthermore, the integration of additional downstream analytical modules, including beta diversity visualisation, differential abundance testing, and longitudinal analysis, would extend VPHYL beyond format conversion. Community-driven schema definitions for Nanopore classifier outputs would also be a valuable parallel initiative, and VPHYL’s parsing architecture is designed to accommodate such standards as they develop. VPHYL is openly available for the community to use, and researchers can parse, standardise, and analyse their Nanopore microbiome abundance data directly at https://venkat-247.shinyapps.io/VPHYL/. No installation, programming experience, or local R environment is required. VPHYL lowers the technical barrier for this conversion step.

## 7. Conclusion

VPHYL is a web application that converts Emu and EPI2ME wf-16s abundance tables from Nanopore 16S data into phyloseq (RData) and BIOM objects and produces a basic summary report. Output is identical to an independent R script, and a case study in which published wf-16s tables were processed as provided, shows that the conversion is reliable. VPHYL therefore provides an accessible and reproducible way to prepare these tables for downstream analysis.

## Supporting information

The application is publicly available at https://venkat-247.shinyapps.io/VPHYL

## Acknowledgement

The authors acknowledge the support of the Indian Council of Medical Research (ICMR) under the grant IIRPSG-2024-01-00486, and Deepak Raghavan Family Foundation (DRFF) Research Program Endowment Fund (IITD/ADFA/20261512442).

