## Supplementary material for "VPHYL: A Web Application for Converting Emu and EPI2ME wf-16s Nanopore 16S rRNA Abundance Tables into phyloseq and BIOM Objects": The application is publicly available at https://venkat-247.shinyapps.io/VPHYL

### (Supplementary Information)


Venkata Sai Reddy Ramireddy[1], Sarvesh Kumar Srivastava[1, 2, *]

[1]Centre for Biomedical Engineering, Indian Institute of Technology Delhi, New Delhi, India

[2]Department of Biomedical Engineering, All India Institute of Medical Sciences, New Delhi, India

*


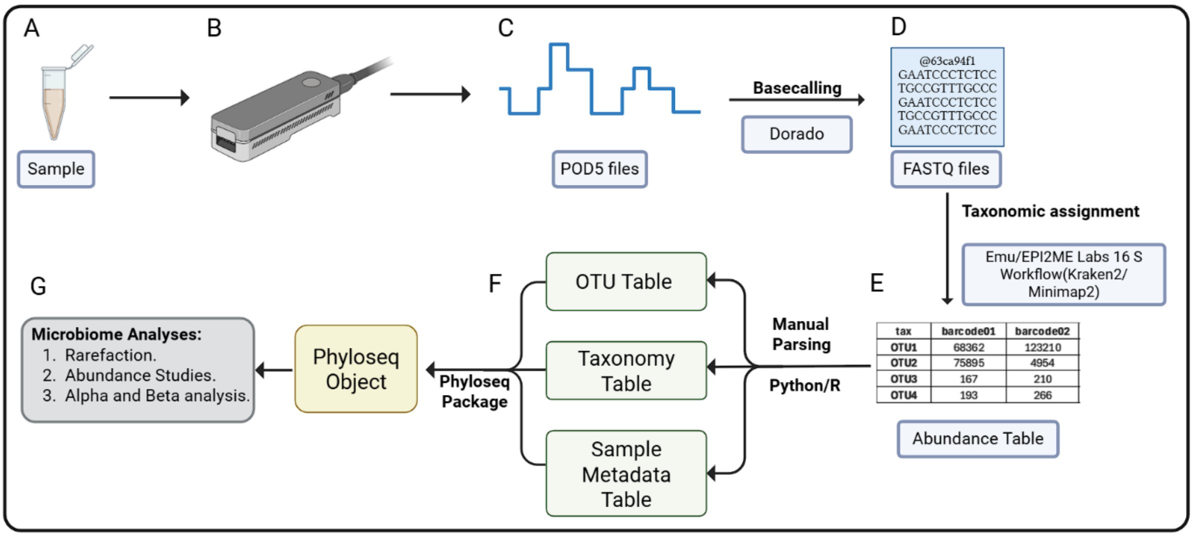


**Supplementary Figure 1**: End-to-End Analytical Workflow for Full-Length 16S rRNA Microbiome Profiling via Oxford Nanopore Technologies. (A) Library Preparation: Systematic extraction of microbial genomic DNA followed by PCR amplification of the full-length (~1,500 bp) 16S rRNA gene (V1–V9 regions). Molecular barcodes are ligated to target ssDNA amplicons, enabling the pooling of diverse samples for multiplexed analysis. (B) Real-time Sequencing: The pooled library is loaded onto a Nanopore flow cell (MinION/GridION/PromethION). (C) Raw Signal Acquisition: ssDNA strands pass through protein nanopores, disrupting the ionic current in a characteristic pattern ("squiggles") recorded as POD5 files. (D) Primary Bioinformatic Processing: Electrical signals are transformed into nucleotide sequences (Basecalling) using neural-network-based algorithms (e.g., Dorado), followed by demultiplexing to sort reads by unique molecular barcodes. (E) Taxonomic Classification: Sequences are converted into taxonomic abundance profiles using specialized profilers with curated reference databases. The EPI2ME 16S workflow utilizes Kraken2 (k-mer based) and Minimap2 (seed-chain-align) with using reference including NCBI_16s_18s, NCBI_16s_18s_28s_ITS, and SILVA_138_1. Simultaneously, the Emu classifier using the Emu_Default (copy-number corrected), RDP_v11.5, or SILVA_v138.1 databases. (F) The Abundances table is used to prepare the required tables like OTU, taxonomy and sample table to merge into a single S4 object using the phyloseq package. G) Downstream Microbiome analysis.

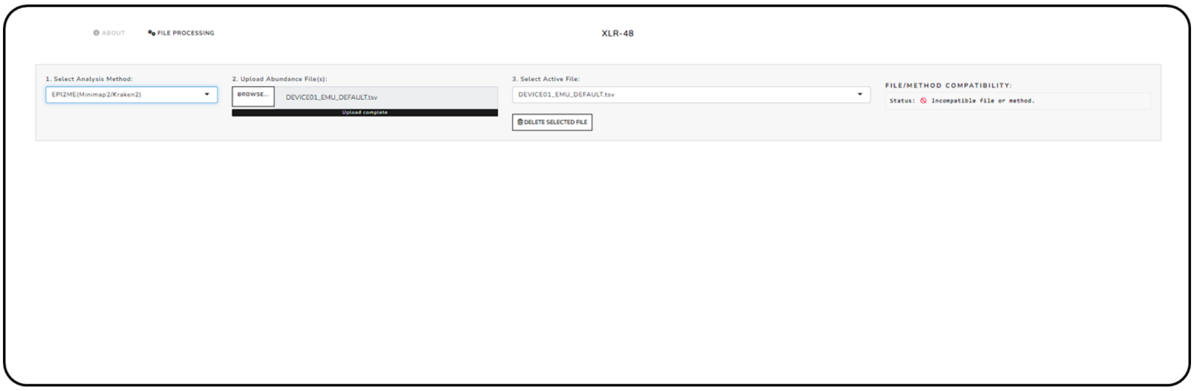


**Supplementary Figure 2:** VPHYL shiny application showing the file compatibility "Incompatible" upon uploading the abundance file processed by "Emu classifier with Emu database" with selecting the incorrect Analysis Method "EPI2ME(Kraken2/Minimap2)

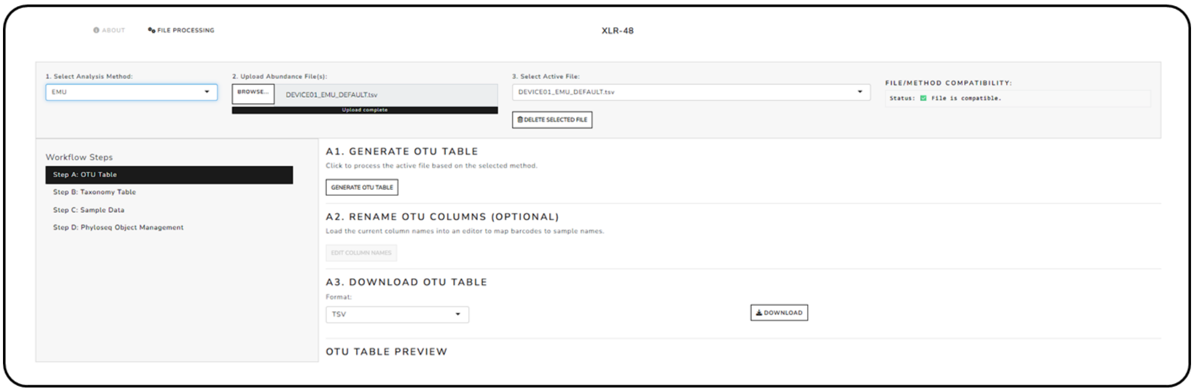


**Supplementary Figure 3:** VPHYL shiny application showing the file compatibility "Compatible" upon uploading the abundance file processed by "Emu classifier with Emu database" with selecting the correct Analysis Method "Emu"

**Supplementary Table 1:** Comparative analysis of VPHYL features versus other microbiome data processing and web-based analysis platforms.

| Tool | Primary Role | User Interface | Metadata Integration | Cross-Classifier Compatibility | Main Outputs | Target Users |
|---|---|---|---|---|---|---|
| **VPHYL** | Standardization, metadata integration, and phyloseq generation | Web GUI | Yes | Emu, Kraken2, Minimap2 | phyloseq, BIOM, CSV/TSV, HTML | Novice (requires prior classification) |
| **TAXPASTA** | Taxonomic profile standardization | CLI | No | Multiple profilers | TSV/CSV, BIOM | Advanced |
| **KrakenTools / kraken-biom** | Kraken ecosystem conversion and formatting | CLI | No | Kraken only | BIOM | Advanced |
| **Pavian** | Interactive exploration and visualization | Web GUI | No | Kraken, Centrifuge, MetaPhlAn | HTML visualizations | Novice |
| **MicrobiomeAnalyst** | End-to-end processing and statistical analysis | Web GUI | Limited (external metadata required) | Classifier-agnostic | Statistical reports and plots | Novice |
| **epi2me2r** | Import of EPI2ME WIMP and AMR results into R | R package | Yes | EPI2ME WIMP and AMR only | phyloseq, metagenomeSeq | Advanced |

**Supplementary Table 2:** The Alpha indices (Observed, Chao, Shannon, and Simpson) were generated using the raw abundance file provided in the paper by Macip, Guillem, et al. [32]. This data was processed using the VPHYL pipeline. The resulting Alpha indices values were compiled into a table, which includes data from **porcine** respiratory microbiome samples collected over different sequencing durations.

| Sequencing Time | ID | barcode01 | barcode02 | barcode03 | barcode04 | barcode05 |
|---|---|---|---|---|---|---|
| 12 | Observed | 16 | 68 | 29 | 80 | 71 |
| | Choa1 | 21 | 76.67 | 35 | 97.5 | 90.43 |
| | Shannon | 1.014 | 1.762 | 1.367 | 0.88 | 1.655 |
| | Simspon | 0.544 | 0.755 | 0.694 | 0.42 | 0.66 |
| 18 | Observed | 17 | 73 | 30 | 83 | 81 |
| | Choa1 | 20.333 | 90.5 | 41.25 | 94.4 | 100 |
| | Shannon | 1.017 | 1.759 | 1.357 | 0.88 | 1.658 |
| | Simspon | 0.545 | 0.754 | 0.692 | 0.42 | 0.661 |
| 24 | Observed | 18 | 75 | 30 | 85 | 83 |
| | Choa1 | 21.333 | 95 | 45 | 100 | 96.6 |
| | Shannon | 1.02 | 1.756 | 1.348 | 0.89 | 1.652 |
| | Simspon | 0.547 | 0.754 | 0.692 | 0.43 | 0.66 |
| 72 | Observed | 22 | 81 | 31 | 89 | 86 |
| | Choa1 | 23.667 | 115.2 | 36.6 | 95.6 | 94.75 |
| | Shannon | 1.035 | 1.743 | 1.346 | 0.9 | 1.642 |
| | Simspon | 0.558 | 0.751 | 0.692 | 0.44 | 0.663 |
| **Sequencing Time** | **ID** | **barcode06** | **barcode07** | **barcode08** | **barcode09** | **barcode12** |
| 12 | Observed | 71 | 76 | 53 | 54 | 71 |
| | Choa1 | 94.33 | 104.11 | 71.333 | 64.11 | 83 |
| | Shannon | 1.78 | 1.696 | 1.601 | 0.963 | 2.33 |
| | Simspon | 0.717 | 0.746 | 0.7 | 0.452 | 0.82 |
| 18 | Observed | 77 | 79 | 56 | 58 | 80 |
| | Choa1 | 84.56 | 105.25 | 67.25 | 68.11 | 110 |
| | Shannon | 1.776 | 1.698 | 1.586 | 0.965 | 2.32 |
| | Simspon | 0.716 | 0.747 | 0.697 | 0.454 | 0.82 |
| 24 | Observed | 77 | 81 | 57 | 61 | 83 |
| | Choa1 | 82.83 | 102.11 | 64.2 | 76 | 93.5 |
| | Shannon | 1.775 | 1.695 | 1.58 | 0.967 | 2.32 |
| | Simspon | 0.717 | 0.747 | 0.696 | 0.457 | 0.82 |
| 72 | Observed | 82 | 84 | 61 | 70 | 88 |
| | Choa1 | 96.25 | 96.75 | 74.2 | 93.33 | 104 |

| | | | | | | |
|---|---|---|---|---|---|---|
| | **Shannon** | 1.774 | 1.689 | 1.579 | 0.975 | 2.29 |
| | **Simspon** | 0.72 | 0.748 | 0.698 | 0.467 | 0.81 |

| Sequencing Time | ID | barcode01 | barcode02 | barcode03 | barcode04 | barcode05 | barcode06 | barcode07 | barcode08 | barcode09 | barcode12 |
|---|---|---|---|---|---|---|---|---|---|---|---|
| 12 | Observed | 16 | 68 | 29 | 80 | 71 | 71 | 76 | 53 | 54 | 71 |
| | Choa1 | 21 | 76.67 | 35 | 97.5 | 90.43 | 94.33 | 104.11 | 71.333 | 64.11 | 83 |
| | Shannon | 1.014 | 1.762 | 1.367 | 0.88 | 1.655 | 1.78 | 1.696 | 1.601 | 0.963 | 2.33 |
| | Simspon | 0.544 | 0.755 | 0.694 | 0.42 | 0.66 | 0.717 | 0.746 | 0.7 | 0.452 | 0.82 |
| 18 | Observed | 17 | 73 | 30 | 83 | 81 | 77 | 79 | 56 | 58 | 80 |
| | Choa1 | 20.333 | 90.5 | 41.25 | 94.4 | 100 | 84.56 | 105.25 | 67.25 | 68.11 | 110 |
| | Shannon | 1.017 | 1.759 | 1.357 | 0.88 | 1.658 | 1.776 | 1.698 | 1.586 | 0.965 | 2.32 |
| | Simspon | 0.545 | 0.754 | 0.692 | 0.42 | 0.661 | 0.716 | 0.747 | 0.697 | 0.454 | 0.82 |
| 24 | Observed | 18 | 75 | 30 | 85 | 83 | 77 | 81 | 57 | 61 | 83 |
| | Choa1 | 21.333 | 95 | 45 | 100 | 96.6 | 82.83 | 102.11 | 64.2 | 76 | 93.5 |
| | Shannon | 1.02 | 1.756 | 1.348 | 0.89 | 1.652 | 1.775 | 1.695 | 1.58 | 0.967 | 2.32 |
| | Simspon | 0.547 | 0.754 | 0.692 | 0.43 | 0.66 | 0.717 | 0.747 | 0.696 | 0.457 | 0.82 |
| 72 | Observed | 22 | 81 | 31 | 89 | 86 | 82 | 84 | 61 | 70 | 88 |
| | Choa1 | 23.667 | 115.2 | 36.6 | 95.6 | 94.75 | 96.25 | 96.75 | 74.2 | 93.33 | 104 |
| | Shannon | 1.035 | 1.743 | 1.346 | 0.9 | 1.642 | 1.774 | 1.689 | 1.579 | 0.975 | 2.29 |
| | Simspon | 0.558 | 0.751 | 0.692 | 0.44 | 0.663 | 0.72 | 0.748 | 0.698 | 0.467 | 0.81 |

**Supplementary Table 3:** The Alpha indices (Observed, Chao, Shannon, and Simpson) were generated using the raw abundance file provided in the paper by Macip, Guillem, et al. [32]. This data was processed using the VPHYL pipeline. The resulting Alpha indices values were compiled into a table, which includes data from **human** respiratory microbiome samples collected over different sequencing durations.

| Sequencing Time | 12 | | | | 18 | | | |
|---|---|---|---|---|---|---|---|---|
| ID | Observed | Choa1 | Shannon | Simspon | Observed | Choa1 | Shannon | Simspon |
| barcode01 | 4 | 5 | 0.351 | 0.157 | 5 | 8 | 0.323 | 0.132 |
| barcode02 | 39 | 40 | 2.744 | 0.891 | 42 | 42.429 | 2.74 | 0.888 |
| barcode03 | 44 | 49 | 2.285 | 0.831 | 47 | 49 | 2.29 | 0.831 |
| barcode04 | 28 | 30.5 | 2.129 | 0.801 | 30 | 30.857 | 2.129 | 0.799 |
| barcode05 | 4 | 4.333 | 1.33 | 0.722 | 4 | 4.333 | 1.33 | 0.722 |
| barcode06 | 10 | 10 | 0.551 | 0.198 | 12 | 13 | 0.528 | 0.186 |
| barcode07 | 8 | 8 | 1.863 | 0.823 | 15 | 16.667 | 2.269 | 0.863 |
| barcode08 | 11 | 12.5 | 2.057 | 0.834 | 13 | 18 | 2.033 | 0.815 |
| barcode09 | 15 | 21 | 2.072 | 0.84 | 16 | 17.5 | 2.161 | 0.852 |
| barcode10 | 5 | 5.5 | 1.129 | 0.56 | 6 | 6.333 | 1.3 | 0.612 |
| barcode11 | 45 | 55.5 | 2.641 | 0.887 | 49 | 58 | 2.655 | 0.888 |
| barcode12 | 14 | 16.5 | 1.158 | 0.54 | 15 | 22.5 | 1.171 | 0.547 |
| barcode13 | 13 | 13.75 | 1.351 | 0.583 | 13 | 16 | 1.343 | 0.576 |
| barcode14 | 17 | 18 | 1.468 | 0.608 | 19 | 19.6 | 1.469 | 0.611 |
| barcode15 | 12 | 12.5 | 0.574 | 0.212 | 13 | 16 | 0.565 | 0.205 |
| barcode16 | 25 | 27.5 | 2.121 | 0.772 | 26 | 29.333 | 2.108 | 0.768 |
| barcode17 | 16 | 23.5 | 1.643 | 0.649 | 18 | 21.333 | 1.695 | 0.666 |
| barcode18 | 11 | 12.5 | 0.102 | 0.035 | 12 | 13 | 0.108 | 0.037 |
| barcode19 | 21 | 22.5 | 0.696 | 0.26 | 21 | 21.333 | 0.684 | 0.258 |
| barcode20 | 9 | 9 | 0.749 | 0.333 | 10 | 10 | 0.747 | 0.325 |
| barcode21 | 53 | 55.143 | 0.804 | 0.307 | 56 | 61.6 | 0.812 | 0.311 |
| barcode22 | 47 | 51 | 1.278 | 0.542 | 50 | 56 | 1.258 | 0.535 |
| barcode23 | 46 | 48.333 | 1.769 | 0.662 | 49 | 54.143 | 1.759 | 0.657 |
| barcode24 | 41 | 48.2 | 1.83 | 0.747 | 43 | 46 | 1.846 | 0.751 |
| Sequencing Time | 24 | | | | 72 | | | |
| ID | Observed | Choa1 | Shannon | Simspon | Observed | Choa1 | Shannon | Simspon |
| barcode01 | 5 | 8 | 0.314 | 0.13 | 5 | 5.5 | 0.337 | 0.143 |
| barcode02 | 44 | 44.5 | 2.742 | 0.886 | 46 | 48 | 2.749 | 0.886 |
| barcode03 | 48 | 48.6 | 2.281 | 0.828 | 49 | 49.2 | 2.28 | 0.829 |
| barcode04 | 30 | 30.429 | 2.128 | 0.801 | 31 | 32.5 | 2.129 | 0.803 |
| barcode05 | 4 | 4.333 | 1.33 | 0.722 | 4 | 4.5 | 1.277 | 0.694 |
| barcode06 | 12 | 13 | 0.51 | 0.18 | 13 | 16 | 0.546 | 0.193 |
| barcode07 | 15 | 17 | 2.242 | 0.859 | 17 | 17.857 | 2.312 | 0.865 |

| | | | | | | | | |
|---|---|---|---|---|---|---|---|---|
| **barcode08** | 14 | 21.5 | 2.088 | 0.827 | 16 | 23.5 | 2.211 | 0.839 |
| **barcode09** | 16 | 16.333 | 2.129 | 0.843 | 16 | 17 | 2.109 | 0.839 |
| **barcode10** | 7 | 7.25 | 1.509 | 0.7 | 7 | 7.333 | 1.539 | 0.718 |
| **barcode11** | 49 | 56 | 2.662 | 0.887 | 51 | 52.667 | 2.66 | 0.887 |
| **barcode12** | 15 | 16.5 | 1.183 | 0.551 | 16 | 18.5 | 1.14 | 0.532 |
| **barcode13** | 14 | 17 | 1.354 | 0.572 | 15 | 18 | 1.388 | 0.581 |
| **barcode14** | 20 | 20.5 | 1.428 | 0.595 | 22 | 22.5 | 1.446 | 0.592 |
| **barcode15** | 14 | 20 | 0.551 | 0.198 | 15 | 18 | 0.567 | 0.205 |
| **barcode16** | 26 | 29.333 | 2.098 | 0.77 | 27 | 28.5 | 2.071 | 0.762 |
| **barcode17** | 20 | 27.5 | 1.673 | 0.651 | 20 | 22 | 1.77 | 0.684 |
| **barcode18** | 12 | 13 | 0.105 | 0.036 | 14 | 16 | 0.104 | 0.035 |
| **barcode19** | 21 | 21 | 0.679 | 0.257 | 21 | 21 | 0.684 | 0.26 |
| **barcode20** | 10 | 10 | 0.749 | 0.324 | 10 | 10 | 0.758 | 0.326 |
| **barcode21** | 56 | 63 | 0.801 | 0.307 | 56 | 59.75 | 0.807 | 0.31 |
| **barcode22** | 50 | 52.625 | 1.262 | 0.535 | 51 | 52.667 | 1.253 | 0.534 |
| **barcode23** | 49 | 52.5 | 1.754 | 0.655 | 50 | 59.333 | 1.753 | 0.654 |
| **barcode24** | 45 | 49 | 1.845 | 0.751 | 49 | 52.5 | 1.848 | 0.752 |

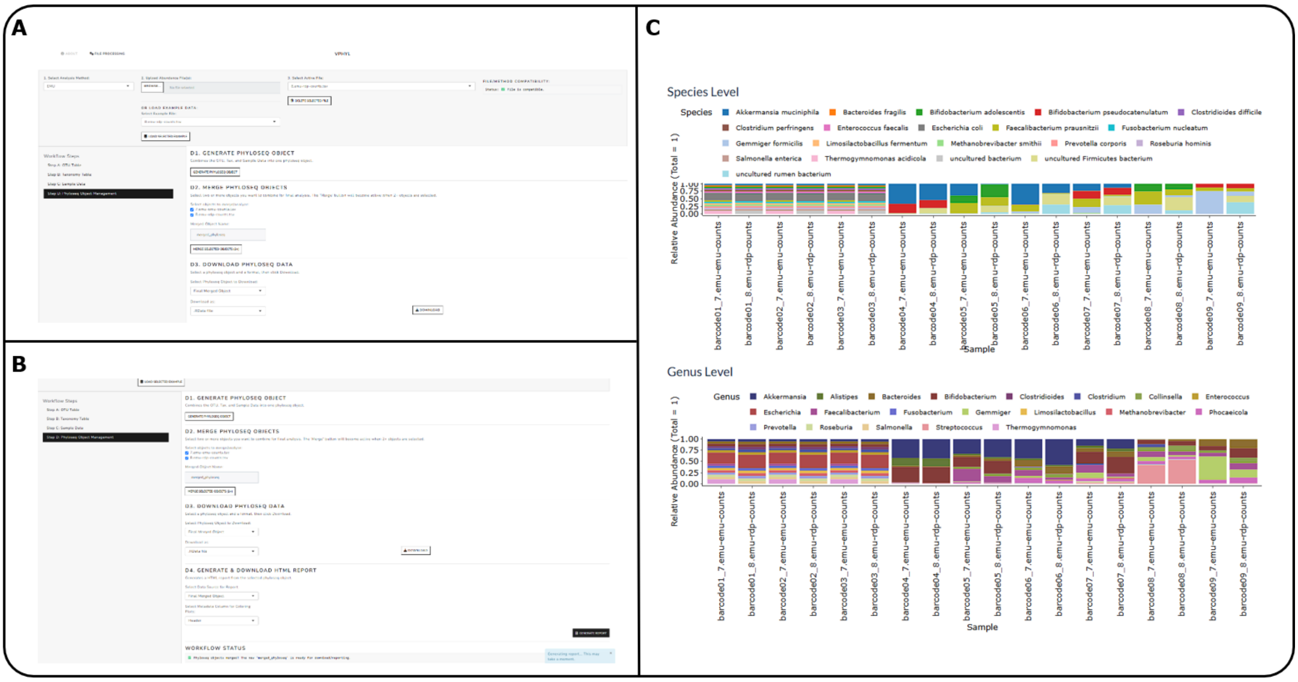


**Supplementary Figure 4**: Demonstration of VPHYL's Multi-File Aggregation and Standardization Across Heterogeneous Classifier Outputs. (A) The application interface demonstrates the independent ingestion and standardization of two disparate raw abundance files from the example dataset (File 1: Emu classifier mapped against the Emu default database; File 2: Emu classifier mapped against the RDP database). Following independent parsing into standardized S4 objects, the components are aggregated into a single, unified dataset ( merged_phyloseq ) using the 'Merge Selected Objects' function in the Step D panel. (B) The interface demonstrates the generation of a unified HTML analytical report directly from the aggregated merged_phyloseq object. (C) Automated compositional visualization at the genus and species levels was generated from the merged dataset. The x-axis labels preserve the classifier-database origin for each barcode, showing VPHYL's capability to seamlessly ingest, combine, and co-visualize samples processed through different workflows within a single downstream analysis.
